# Extensive OMICS resource for Sf21 and Tni cell lines

**DOI:** 10.1101/2021.04.06.438574

**Authors:** Bence Galik, Jonathan J.M. Landry, Joanna M. Kirkpatrick, Markus Hsi-Yang Fritz, Bianka Baying, Jonathon Blake, Bettina Haase, Paul G. Collier, Rajna Hercog, Dinko Pavlinic, Peggy Stolt-Bergner, Hüseyin Besir, Kim Remans, Attila Gyenesei, Vladimir Benes

## Abstract

Insect-derived cell lines, from *Spodoptera frugiperda* (Sf21) and from *Trichoplusia ni* (Tni), are the two most widely used cell lines for recombinant protein expression in combination with the Baculoviral Expression Vector System (BEVS). Genomic sequences and annotations are still incomplete for Sf21 and absent for Tni. In this study, we present an approach using different sequencing data types, including short-read sequencing, long synthetic and Oxford Nanopore reads, to build genomes. The Sf21 and Tni assemblies contain 4,020 scaffolds of 463 Mb in size with N50 of 364 Kb and 2,954 scaffolds of 332 Mb in size with N50 of 326 Kb, respectively. Furthermore, we built a new gene prediction workflow, which integrates transcriptome and proteome information using pre-existing tools. Using this approach, we predicted 21,506 Sf21 and 14,159 Tni genes, generated and integrated proteomic datasets to validate predicted genes and could identify 5577 and 4919 proteins in the Sf21 and Tni cell lines respectively. This integrative approach could be theoretically applied to any uncharacterized genome and result in valuable new resources. With this information available, Sf21 and Tni cells will become even better tools for protein expression and could be used in a wider range of applications, from promoter identification to genome engineering and editing.

## Introduction

Recombinant protein production widely uses baculovirus-mediated expression in insect cells (Jarvis 2009; van Oers 2011). In recent years, many new developments have made these tools safe, efficient, convenient to use and easy to scale up. These cell lines can perform post-translational processing during eukaryotic protein production, be employed for vaccine synthesis and also for adeno-associated virus production designed for gene therapy (van Oers et al. 2015). In these different applications, two cell lines, namely Sf21 (IPLB-Sf21AE) derived from *Spodoptera frugiperda* (Vaughn & Fan 1997), and Tni (BTI-TN-5B1-4) derived from *Trichoplusia ni* tissues (Wickham et al. 1992; Davis et al. 1992) are extensively utilized. Engineering of these cell lines is of great interest to improve protein production. As an example, modifications of the glycosylation pathway in these cells have been subject to many investigations over the past two decades (Xu & Ng 2015; Hollister & Jarvis 2001; Hollister et al. 2002; Breitbach & Jarvis 2001). Increasing the yield of protein synthesis via post-transcriptional gene silencing, mediated by RNA interference technologies, was also achieved by suppressing genes involved in apoptosis, using a DNA vector-based approach combined with endogenous dsRNA expression (Lai et al. 2012).

Deep characterization of genomes has been made possible with the development of extremely rapid sequencing technologies (Heather & Chain 2016). Achieving a better understanding of cell line’s genomes has the potential to unlock the power that would allow specific functionalities of the cells to be engineered; for example, faster and easier adaptation to various culturing conditions or improving the levels of protein production. In addition, gaining access to genomic information would also allow for proteomic identification of native interaction partners or host organism contaminants.

Genomics and transcriptomics information for the Sf21 cell line (Kakumani et al. 2014, 2015), as well as genome (Fu et al. 2018) and transcriptome (Yu et al. 2016) sequences for the Tni cell line have been published.

Here, we present more comprehensive resources for both Tni and Sf21, which include detailed information from not only genomic and transcriptomic, but also proteomic data. We use a multi-sequencing library type approach to assemble genomic and transcriptomic sequences, as well as an innovative annotation workflow to predict genes. In addition to the functional characterization of these gene sets, we also report proteome expression profiles of these two cell lines. These comprehensive resources are deposited in Lepbase (Challis et al. 2016) and are available to the community. The mass spectrometry proteomics data have been deposited in the ProteomeXchange Consortium via the PRIDE (Vizcaíno et al. 2016) partner repository with the dataset identifier PXD010282.

## Results

### Genome assembly

The assembly workflow consists first of a contig assembly using paired-end (PE) reads. Secondly, based on this preliminary assembly, scaffolds were built using mate pair (MP) reads. Then, long-synthetic (LS) reads were employed to construct scaffolds again on the previous assembly. Finally, Oxford Nanopore Technologies (ONT) reads were used to extend the previous assembly to its final state (Figure 1). Figure 2 and Table S1 present the assembly statistics after each of the workflow steps and demonstrate the value of the successive integration of different sequencing data types. Indeed, the N50 (i.e. the sequence length of the shortest contig at 50% of the total genome length) and the length of the largest DNA sequence assembled show an significant increase throughout the assembly process for both genomes.

**Figure 1.**
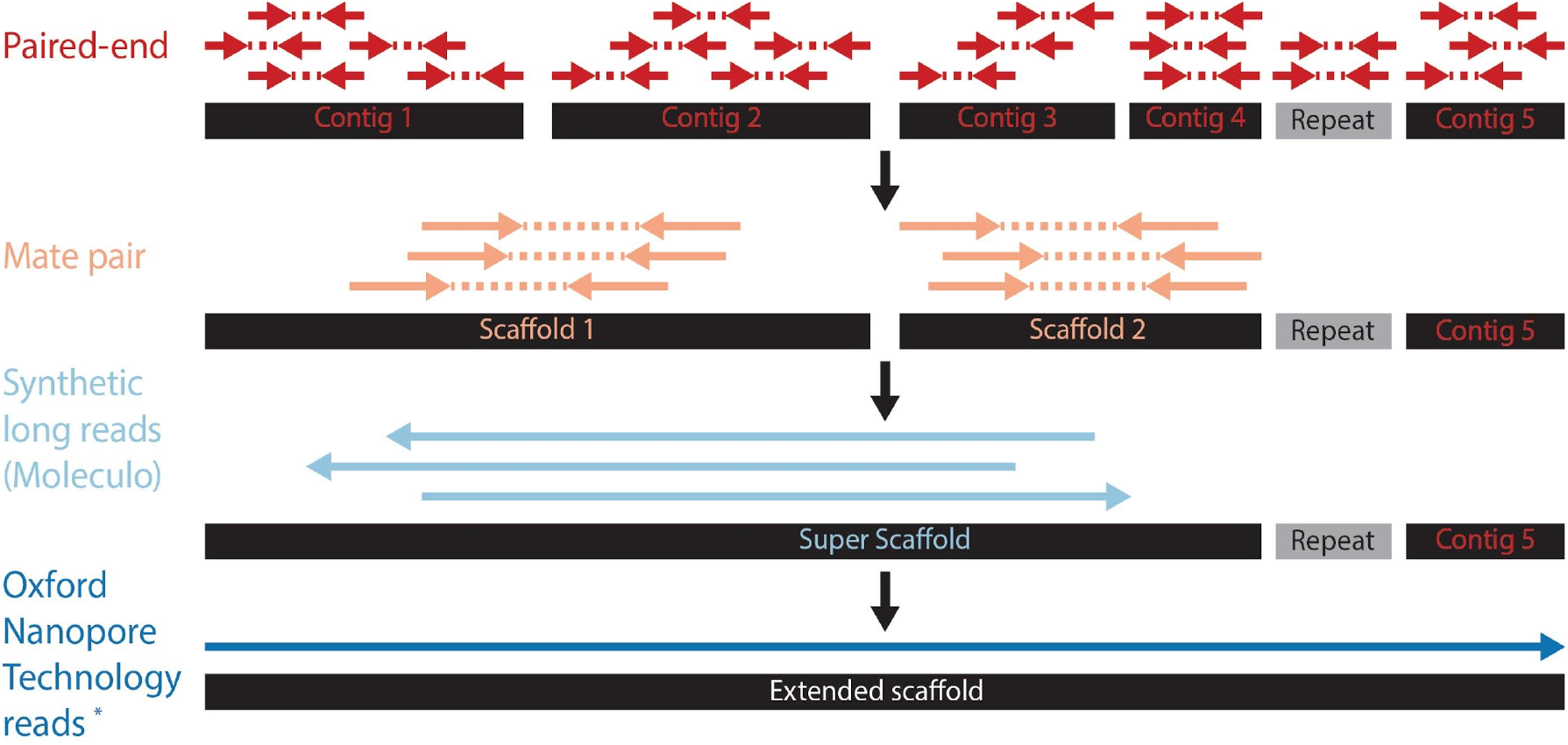
Genome assembly workflow. Schematic representation of the hybrid assembly approach used to build the genome of both cell lines. First, the paired-end reads are used to build contigs. Second, mate pair data are used to scaffold the contigs. Third, synthetic long reads are used construct super scaffolds. Finally, ONT reads extend the super scaffolds to the final genome assembled sequences provided with this paper.

**Figure 2.**
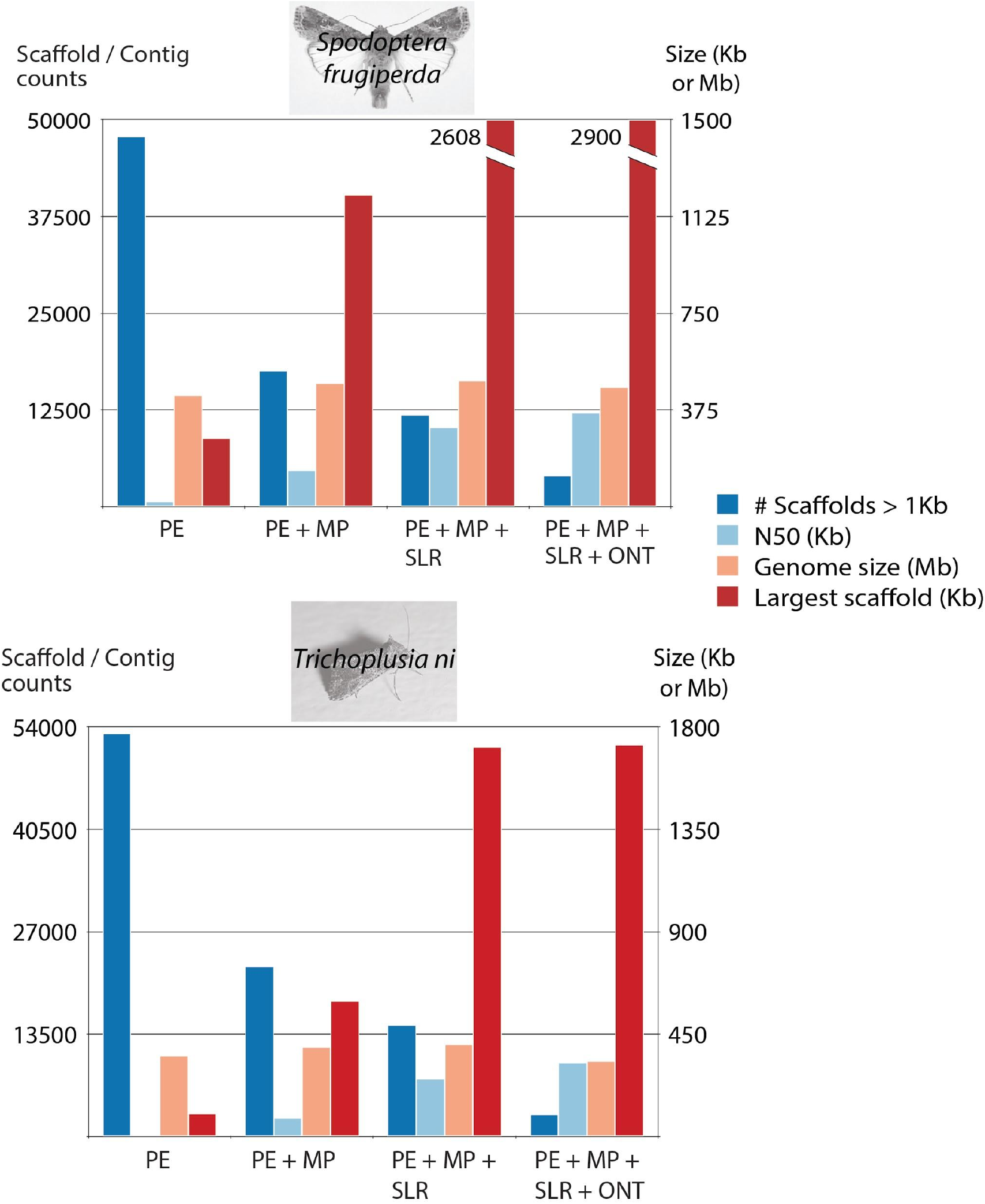
Genome assembly metrics improvement with the integration of the different data types. Genome assemblies metrics, including the number of scaffolds (larger than 1Kb in size), the N50 (Kb), the genome size (Mb) and the largest assembled scaffold (Kb) for Sf21 (top panel) and Tni (bottom panel) at each stage of the workflow, with only paired-end (PE) data, and then integrating mate pair (MP), synthetic long read (SLR) and finally Oxford Nanopore Technology (ONT) datasets.

It is interesting to notice the decrease of the scaffolds number larger than 1Kb after the integration of the ONT reads, for Tni and Sf21 cell lines (Figure 2). These reads, longer than the Illumina datasets, shows their impact on the assembly process and notably also on the N50 value. The current development and progress in terms of throughput and stability of ONT sequencing represent an exciting time for new assembly projects using this technology. With this approach, we were able to finally assemble 4,020 Sf21 scaffolds (equal or larger than 1 Kb), which represent a total size of 463 Mb with the N50 of 364 Kb (Table 1). Our new assembly outcompetes the first Sf21 genome assembly published by Kakumani et al. in 2014, which reported 37,243 scaffolds of size 358 Mb with N50 of 53.7 Kb. Gouin et al. (2017) used a different approach on the “C” and “R” *Spodoptera frugiperda* strains to assemble 4,222 joined, plus 11,628 singleton scaffolds, with a final N50 of 144 Kb. The genome size is 438 Mb and 371 Mb for the “C” and the “R” strain, respectively.

**Table 1:** Short summary of Sf21 and Tni *de novo* genome assemblies. The N50 value is defined as the sequence length of the shortest contig at 50% of the total genome length. L50 value represent the smallest number of contigs to which their cumulative length represents half of genome size.

|  | Sf21 assembly | Tni assembly |
| --- | --- | --- |
| # sequences ($\geq 1$ Kb) | 4,020 | 2,954 |
| Largest scaffold (bp) | 2,900,257 | 1,722,548 |
| Total length (bp) | 463,041,686 | 332,103,479 |
| N50 (bp) | 364,523 | 326,309 |
| L50 | 315 | 293 |
| GC (%) | 38.42 | 36.30 |
| # N's per 100 Kb | 681.43 | 499.33 |

In addition, our assembly compares well with the genome size of one of the closest-related species, i.e. *Bombyx mori* (432 Mb) (Consortium 2008).

Published *Spodoptera frugiperda* genome sequences (Kakumani et al. 2014; Gouin et al. 2017) were compared to our assembled Sf21 scaffolds. Sequence similarity between ours and Kakumani et al. (2014) as Gouin et al. (2017) is shown in Figure 3A and 3B, respectively. On the two dot plots, the overall sequence similarity is high (i.e. red diagonal line), even though the genome sizes are different. The shorter scaffolds presented in the first paper (Kakumani et al. 2014) are depicted with small dots and short lines at the top of Figure 3A. These shorter scaffolds might include short similarities (repeats) between the two genomes and therefore linked to each other’s in this cloud of points.

**Figure 3.**
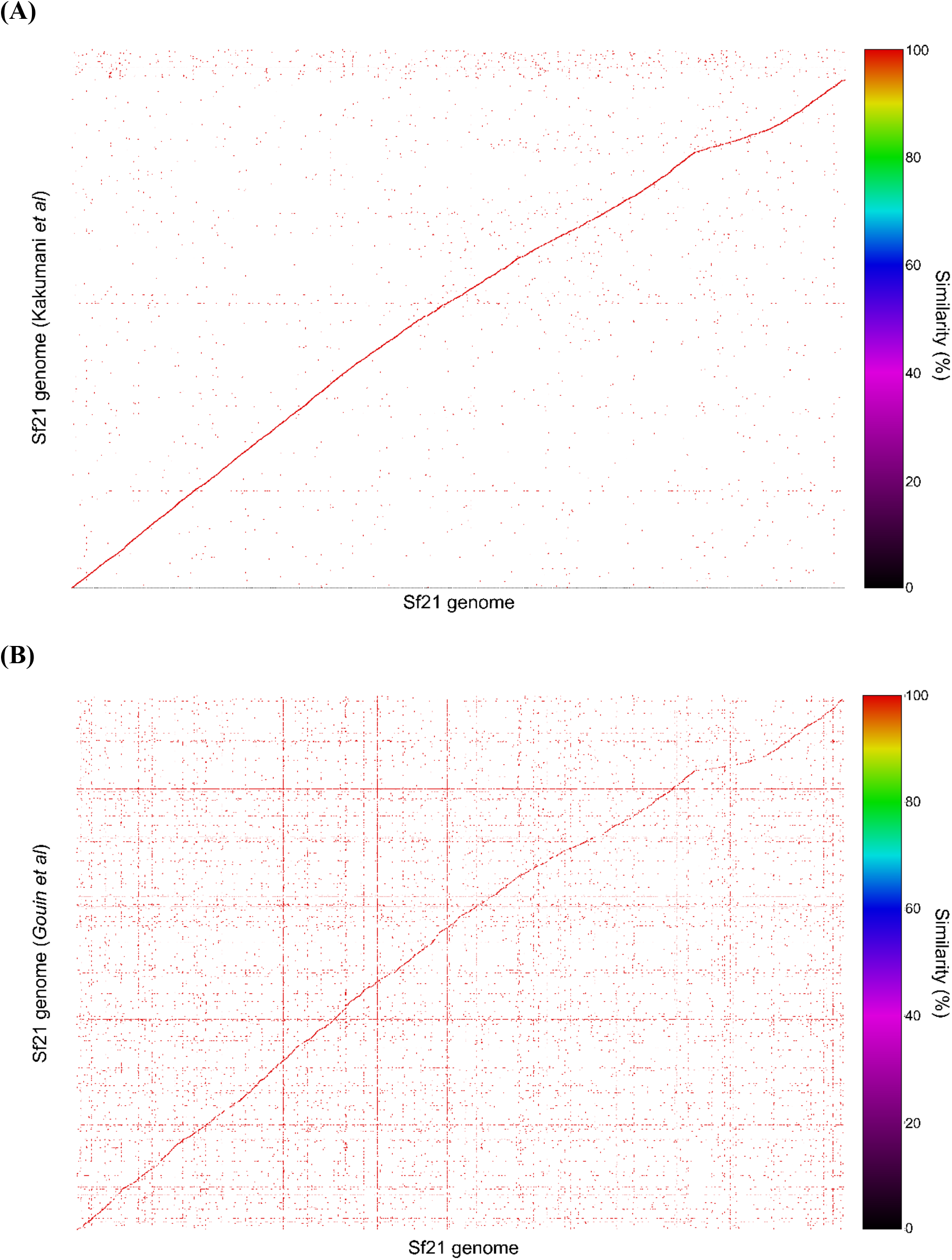
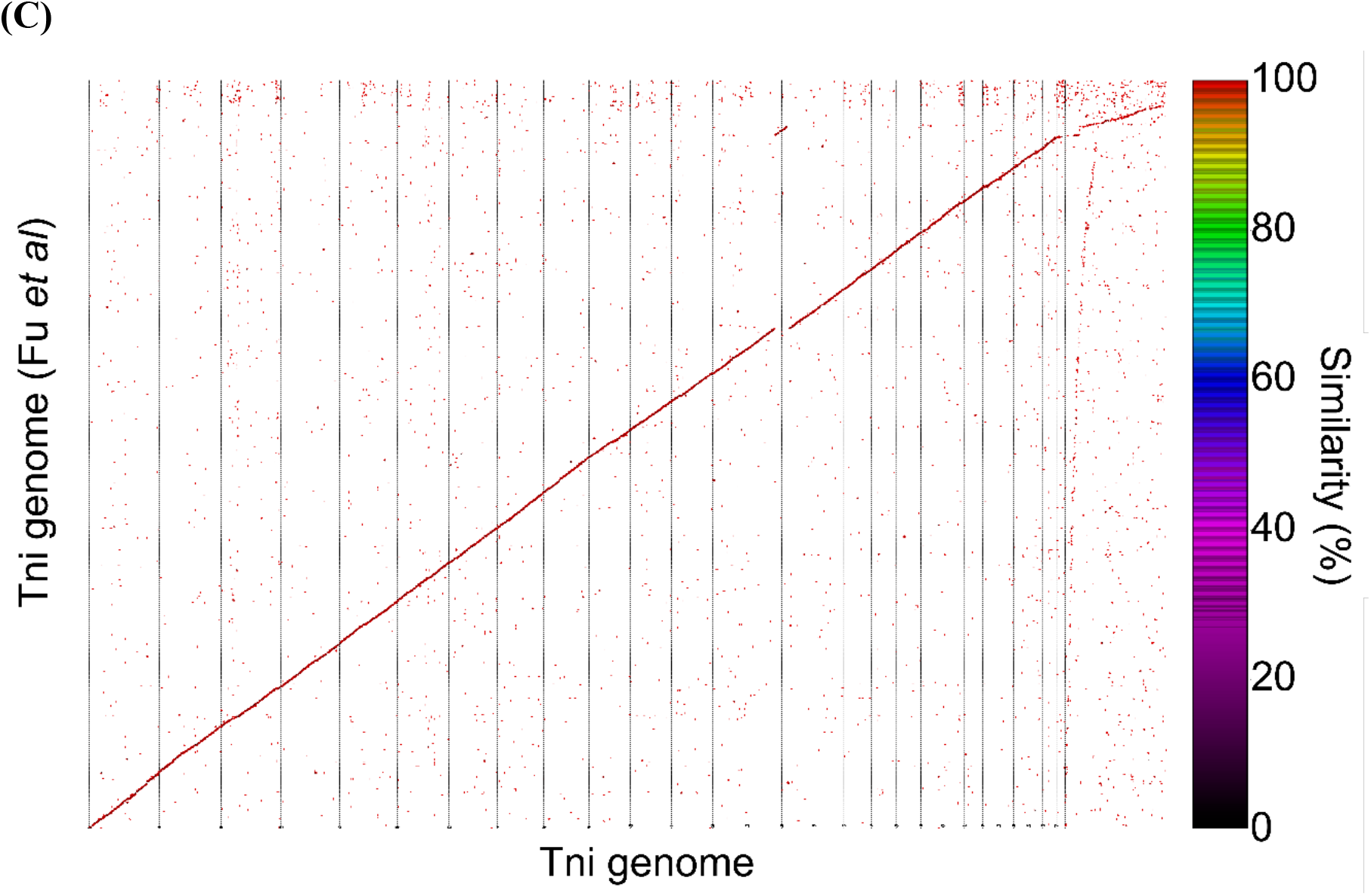
Comparison of the published and our Sf21 and Tni genome assemblies. Alignment dot plots are generated by Mummer (Kurtz et al. 2004) and show the sequence similarity of the 2 genomes. The plot is comparing (A) the Sf21 genome published by Kakumani et al. in 2014 and our assembled Sf21 genome; (B) the Sf21 genome published by Gouin et al. in 2017 and our assembled Sf21 genome; (C) Tni genome published by Fu et al. in 2018 and our assembled Tni genome. For each comparison, the two genome sequences are ranked by size and plotted independently on the x- and y-axis. When two sequences are mapping to each other a colored-line or a dot is plotted. Forward matches are colored in red and reverse matches in green. If the 2 sequences are perfectly the same a red line from bottom left to top right would be drawn. Dots above or below the main slope are indicating translocations or repeats.

Figure 3B compares the latest Sf21 genome (Gouin et al. 2017) and shows more background points and lines compared to Figure 3A, which, counterintuitively, is a sign of better genome assembly completeness. In Figure 3B, scaffolds containing mostly repeated regions, visualized at the top of Figure 3A, are not present. Instead, these repeats are distributed and assembled across both genomes, resulting to a high degree of connectivity (i.e. horizontal or vertical lines and points) between these similar regions on both axis.

Our Tni assembly consists of 2,954 scaffolds (equal to or larger than 1 Kb), which represent a total size of 332 Mb with the N50 of 326 Kb (Table 1). The genome size is smaller than Sf21, but the N50 and the number of contigs are in the same range. The newly published Tni genome (Fu et al. 2018) is a more complete assembly, applying an additional Hi-C step for extra scaffolding, to generate 1031 scaffolds (of which more than 90% of the genome assembled into 28 major scaffolds) with N50 of 14.3 Mb and a genome size of 368.2 Mb. The comparison of the aforementioned genome and our work are shown in Figure 3C. The alignment contains only a few gaps and one major rearrangement over one of the largest scaffolds, which indicates a large overlap between our assembly and the published Tni genome. The only gap could be caused by a misassembly or real structural difference.

The average number of reads covering each base of the genome ranges from 94 for Sf21 to 64 for Tni cells. In addition, more than 89% of all reads from the two PE libraries for each cell line, Sf21 and Tni respectively, are mapping to the assemblies (sequences larger than or equal to 10 Kb) and are properly paired. These two indications support the high quality of each base of the built sequences associated with high sequence coverage. The Core Eukaryotic Genes Mapping Approach (CEGMA) was used to identify a set of 248 conserved core eukaryotic genes (CEGs) into the assembled sequences. A large percentage (higher than 81%) of the CEGs were fully found in the Sf21 and Tni genomes respectively and confirm the high degree of completeness of our assemblies. More than 90% of these genes could also be partially identified in the Tni genome, indicating that some parts of the assembly still remain fragmented.

Short long interspread Nuclear Elements (SINEs), long interspread elements (LINEs), long terminal repeats (LTRs), DNA elements, small RNAs, satellites, simple repeats, low complexity regions and unclassified features comprises 25.5% and 16.4% for the Sf21 and Tni genomes, respectively. These numbers compare well with other repetitive sequence proportions in insect genomes; for example, *D. melanogaster* 28.9%, *A. gambiae* 14.1% and the closest related species genome, *Bombyx mori* 22.4% (Osanai-Futahashi et al. 2008) (based on RepeatMasker Genomic datasets). These observations indicate the capability of our genome assembly workflow to better resolve the general repeat pitfall that is associated with standard short-read Illumina assemblies.

### Transcriptome assembly

We assembled the RNA-seq data to identify the general expression profile. In total, 24,992 transcripts with an N50 of 1,855 bp and 41,041 transcripts with an N50 of 1,850 bp and 2,076 were assembled for Sf21 and Tni respectively after Cluster Database at High Identity with Tolerance (CD-HIT) runs (min. 90% similarity). The important difference in terms of assembled transcript number might be explained by the sequencing reads generated (~91M and ~240M reads for Sf21 and Tni respectively). Table S2 summarizes the different transcriptome assembly metrics.

The published datasets reported ~24K assembled transcripts for Sf21 (Kakumani et al. 2015) and ~25K assembled transcripts for Tni (Yu et al. 2016). We report significantly more transcripts for Tni. The difference might be attributed to the higher sequencing depth, the library types (stranded/ non-stranded) and to the harvesting strategy, where we collected for Tni 4 different time points (0h, 4h, 8h and 24h after passage) in order to capture a wider mRNA molecule catalog for this cell line. In addition, we could not exclude that our Tni transcript assembly contains misassemblies or errors.

The numbers of complete, fragmented and missing conserved insect orthologs across species are quantified in our transcript assemblies, as well as in available transcriptome resources (Kakumani et al. 2015; Yu et al. 2016) and the results are reported in Figure 4. The proportion of missing insect orthologs appears to be very similar. Our assemblies also decrease the number of fragmented orthologs. Only the Sf21 transcriptome assembly exhibits higher numbers of completed and non-duplicated orthologs, compared to the previously published datasets. On the contrary, the Tni transcriptome assembly contains slightly fewer complete, and more duplicated orthologs than the published resource.

**Figure 4.**
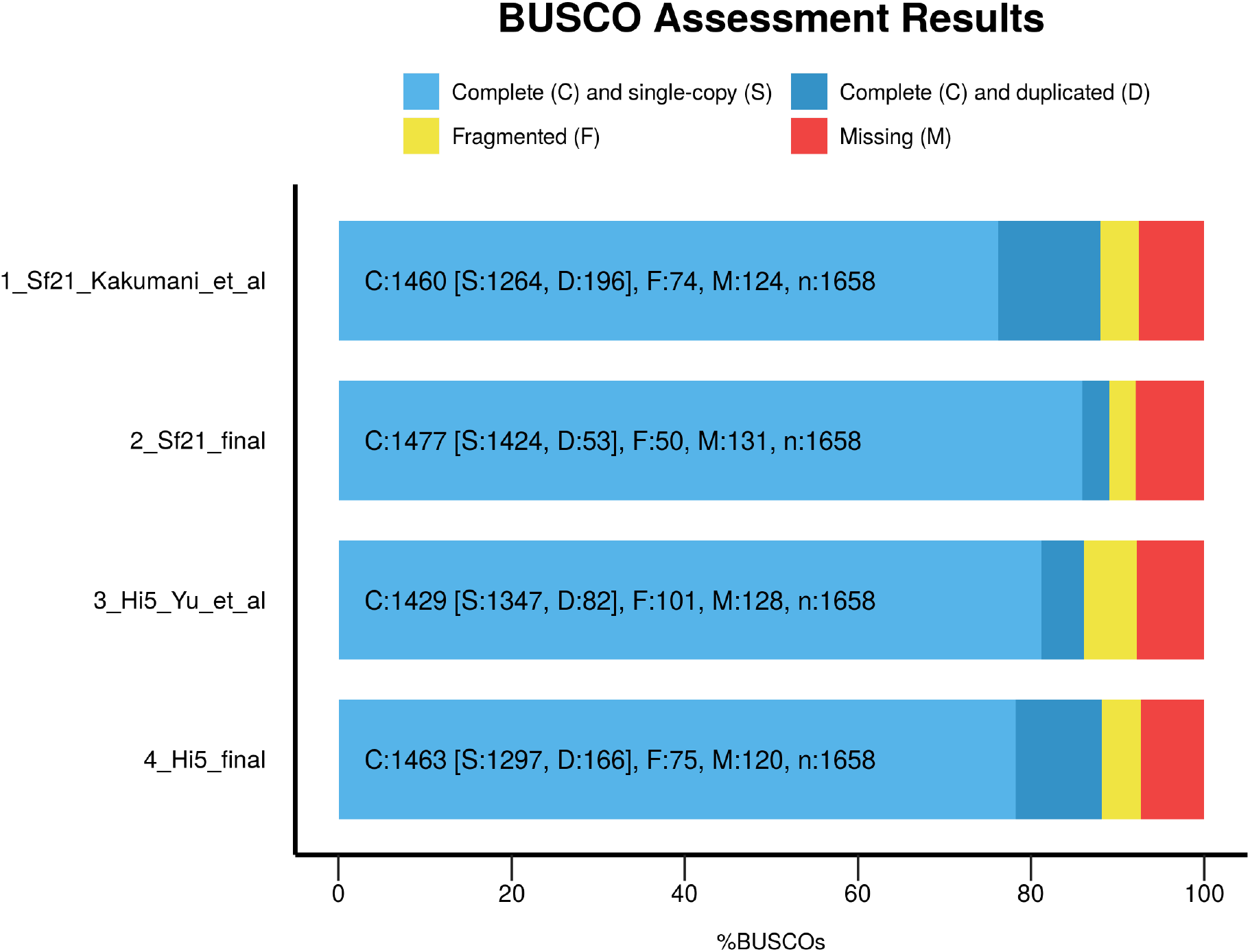
Orthologs distribution for Sf21 and Tni transcriptomes as well as published transcriptomic datasets. The bar charts represents the ortholog distributions found by Benchmarking Universal Single-Copy Orthologs (BUSCO, Simão et al. 2015) for the following categories: complete (C), including single-copy (S) and duplicated (D); fragmented (F); missing (M) across the Sf21 and Tni transcriptomes, generated in this study, as well as the published ones for Sf21 (Kakumani et al. 2015) and High Five (Yu et al. 2016).

In addition, when comparing the published transcript sequences to our newly constructed transcripts, ~ 86% of Sf21 and ~ 55% of Tni sequences were found in our datasets. It is also worth noting that only one contig in each of the available transcriptomes (5881 and 262 for Tni and Sf21 respectively) could not be aligned to any of our sequences.

Transcripts from the different sources were also mapped back to our constructed genomes and the mapping percentage (99% for (Kakumani et al. 2015) and ours) is very high regarding Sf21 and just a little bit lower for Tni (95% and 94% for (Yu et al. 2016) and ours respectively). Finally, to evaluate how complete the assembled transcripts are, they were aligned to the SwissProt database. 3,211 (~ 11.3% for Sf21) and 3,699 (~ 6.4% for Tni) transcripts were reported to be nearly full-length transcripts, when their alignments cover more than 80% of one reference protein. The lower number for the Tni dataset is caused by the higher amount of total assembly transcripts. Apart from this, the main metrics for Sf21 and Tni datasets are quite similar (see Table S2). The same analysis was repeated using the previously published transcripts and 3436 (~ 14.3%) and 3,218 (~ 12.8%) were nearly full-length ones.

Generally, our main objective regarding transcriptome assembly is to use the information to refine the gene prediction accuracy. The preceding comparisons serve to demonstrate that our assemblies are essentially comparable to the published datasets and could be used for the subsequent steps.

### Functional annotation

With our genome annotation workflow, shown in Figure 5, the integration of external evidence, such as the transcript information, allowed us to predict 21,572 Sf21 and 14,594 Tni protein-coding genes. These predictions compare well with the literature. For instance, in Sf21, 11,595 protein-coding genes were predicted using genome assembly (Kakumani et al. 2014) and later 26,390 protein-coding genes using a combination of genome and transcriptome assembly (Kakumani et al. 2015). For Tni, only one publication (Yu et al. 2016) described 13,732 protein-coding genes predicted from transcriptome assembly.

**Figure 5.**
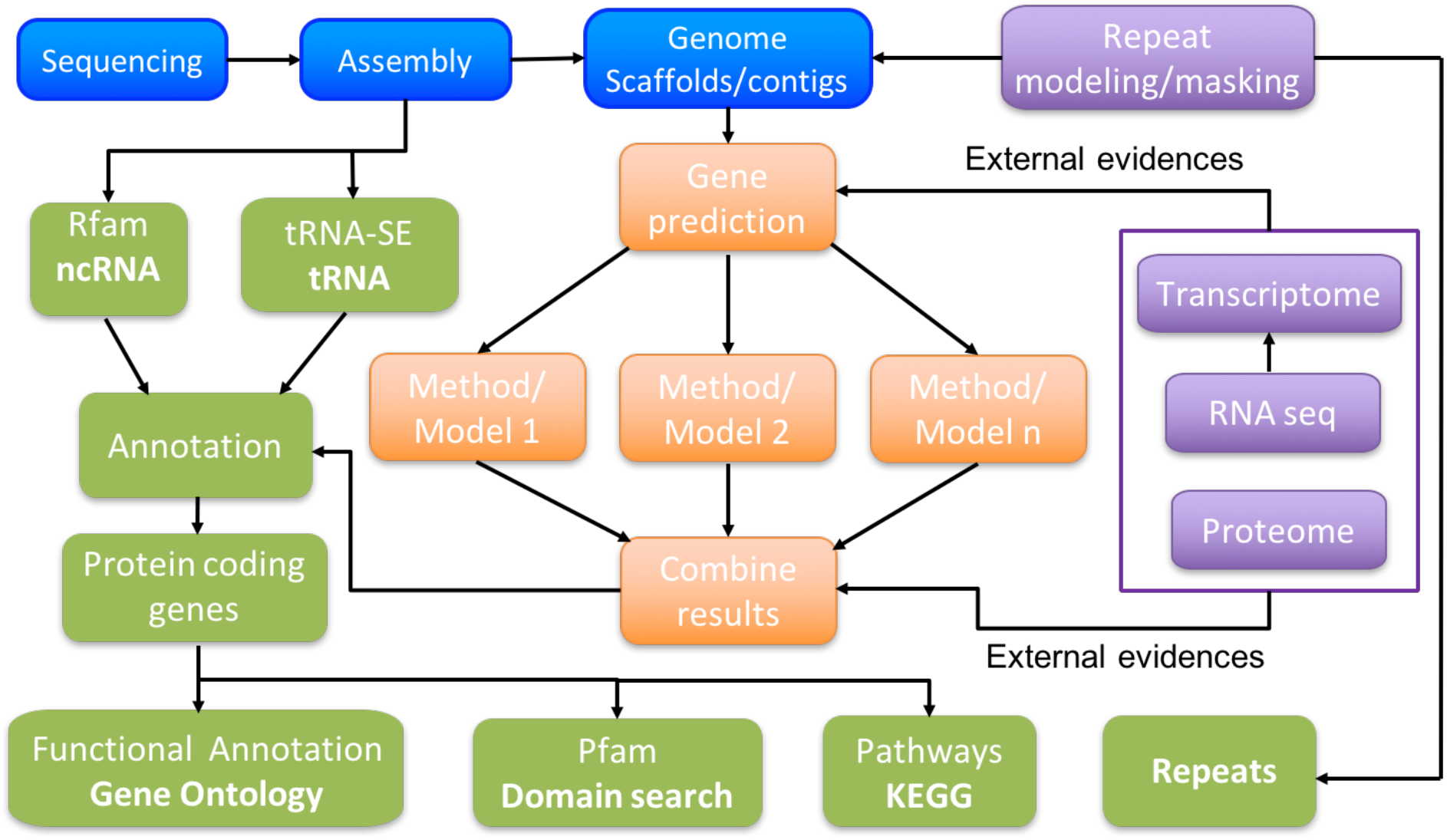
Genome annotation workflow. The workflow chart shows the main steps of our genome annotation method. After masking repeats in the genome assembly, protein-coding genes were predicted using mRNA assembled transcripts as external evidence to support the gene model. The annotation of the predicted genes relies on different databases.

After annotation of the protein-coding genes, Web Gene Ontology Annotation Plot (WEGO, (Ye et al.) was used to visualize general functions for both genomes and proteomes, based on GO terms in the three main categories (Figure 6). Both genomes are very similar. The likely reason for this high similarity is that both cell lines originate from the ovarian cells of their respective organisms. However, although we showed that our genome assembly and annotation methods are quite powerful, we still do not have a chromosome level assembly and there is a chance that certain genes are not captured if they are not active and there is no additional evidence (eg. at the transcript level) to support their identification. This could explain the differences and the missing categories in the WEGO analysis.

**Figure 6.**
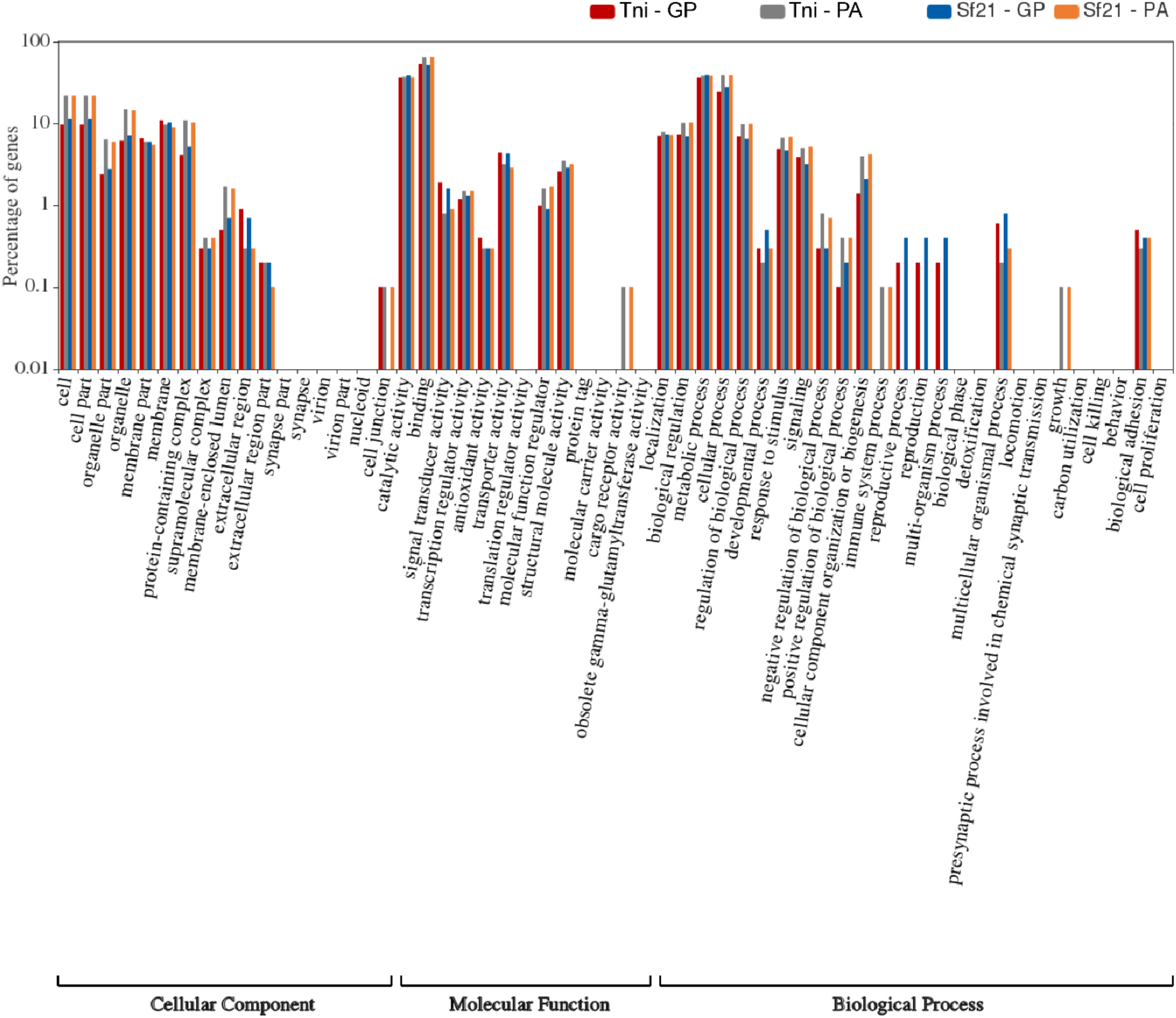
Gene Ontology annotation of Sf21 and Tni protein-coding genes based on the gene prediction and from the protein analysis. It shows the InterProScan GO annotation results of protein-coding genes from Tni and Sf21 based on the hits from the gene prediction (GP) and the protein analysis (PA). The results are summarized in three main GO categories at the second level: Cellular component, Molecular function and Biological process. Right Y-axis is the gene count in that function item, the log(10) scaled left Y-axis is the corresponding percentage of gene number. The plot was obtained using WEGO 2.0 (Ye et al. 2018).

Rfam and tRNAscan-SE tools were used to predict non-protein-coding genes and other features of the genomes. Altogether, 396 Sf21 and 247 Tni ncRNAs, including rRNA, scRNA, miRNA, miscRNA and Cis-reg elements were identified. Moreover, Rfam is also able to predict, among others, tRNAs, too. Here we report only those 1,233 Sf21 and 1,965 Tni tRNAs that were predicted by tRNAscan-SE. Figure 7 and table S3 show the numbers in detail for all different types of genomic features.

**Figure 7.**
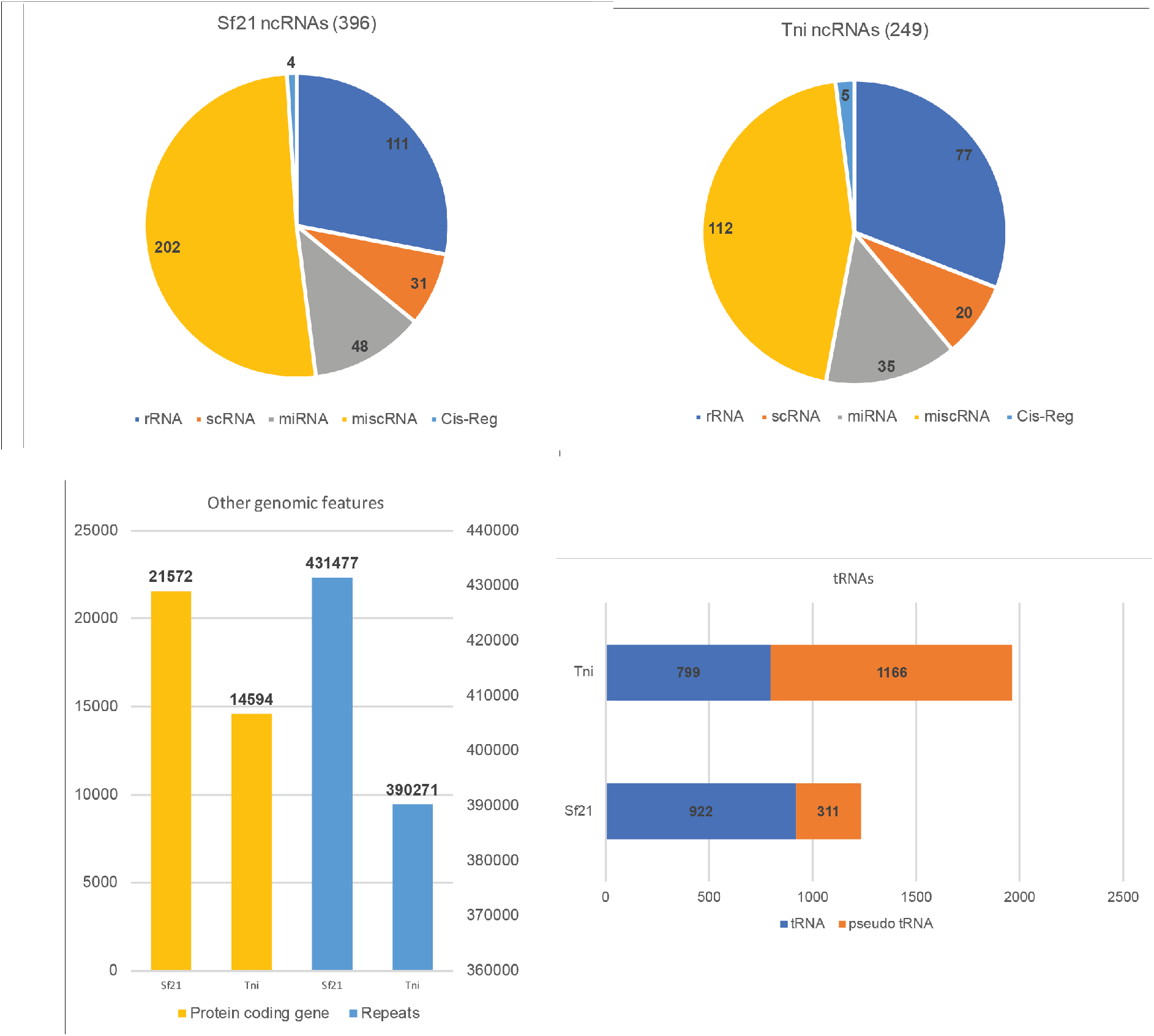
Genomic features. Classification of the gene predictions for SF21 and Tni genomes for (A) ncRNAs, (B) Protein coding genes and repeats, (C) tRNAs.

Using the KAAS portal we assigned Kegg Orthology (KO) IDs to 21,506 and 14,159 Sf21 and Tni protein-coding genes respectively. We focused only on one of the most important pathways in these cell lines, which is that of N-glycosylation (Figure S1). Cytoscape and KEGGREST were used to draw the pathway based on the reference from the KEGG website. The KO IDs and available transcript hit evidence together were used to mark each identified element on the map. Thereby, we had information as to whether an element is only predicted with gene prediction methods, or whether external evidence exists as well. Furthermore, we have the additional layer of information as to whether that gene is expressed or not. All genes involved in the N-glycolsylation pathway could be identified for Sf21, whereas some do not have any evidence for Tni. This might indicate slight differences between *B. mori* and Tni pathways. As an example, the enzyme beta-galactoside alpha-(2,6)-sialyltransferase (EC 2.4.99.1) is present in Tni and absent in *B. mori*. Another possibility could suggest the inability of our assembly and/or gene annotation pipeline to describe some components of the pathway. Some glucosyltransferases (ALG3, ALG6, ALG 7 and ALG10) were, for instance, absent in Tni.

Applying the MaxQuant and the Andromeda tools on the proteome datasets, 5577 Sf21 and 4917 Tni proteins were identified during the analysis respectively. The Log10 transformed iBAQ scores were used to rank and plot the determined proteins from the most abundant to least abundant (Figure 8) The profile of protein abundance reveals a typical S-shaped curve for both Sf21 and Tni covering 5 orders of magnitude of dynamic range, as is expected from a cell line analysis of this type. Also, the WEGO analysis showed the two proteomes are very similar in most of the functional categories (Figure 6).

**Figure 8.**
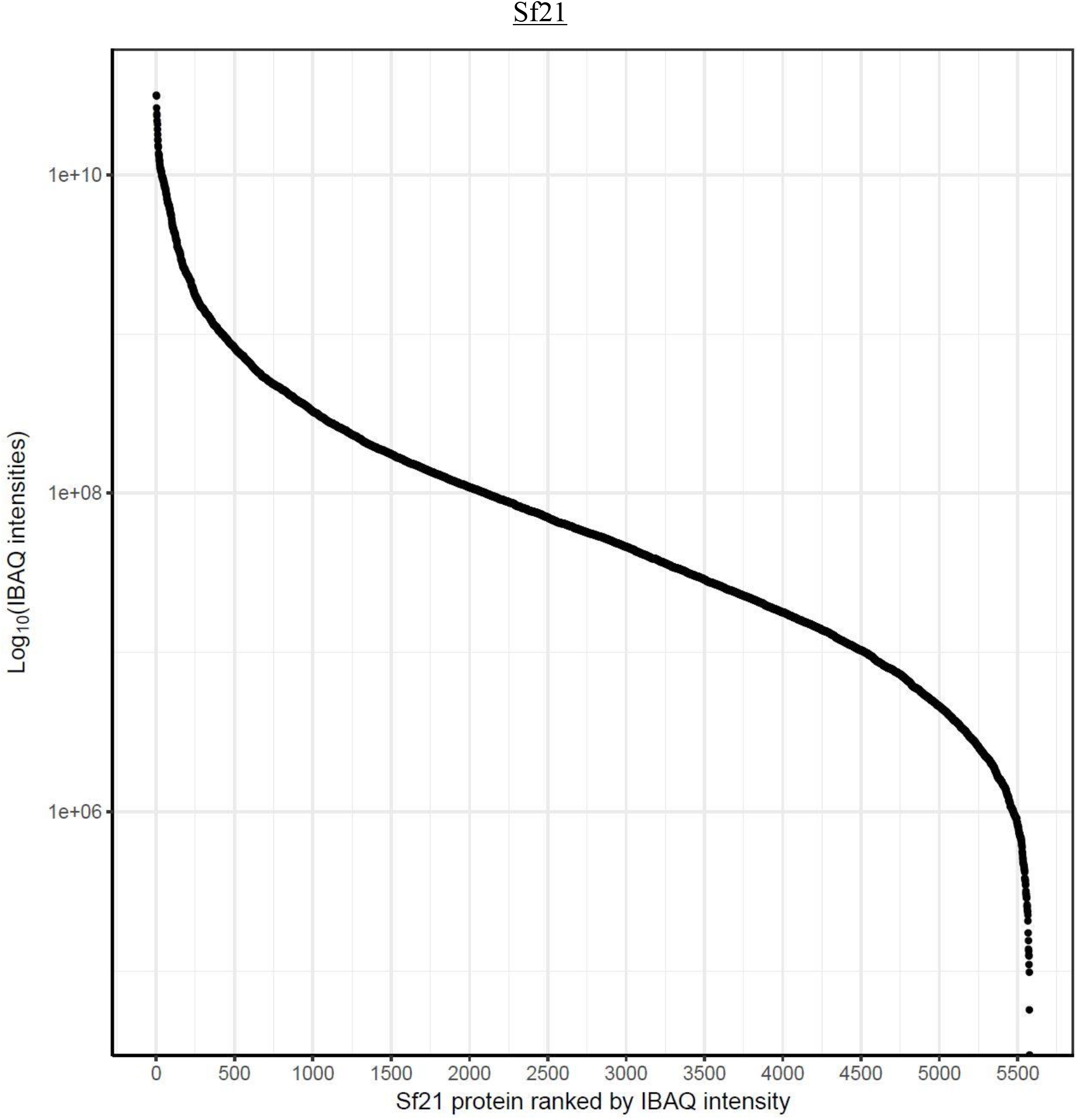
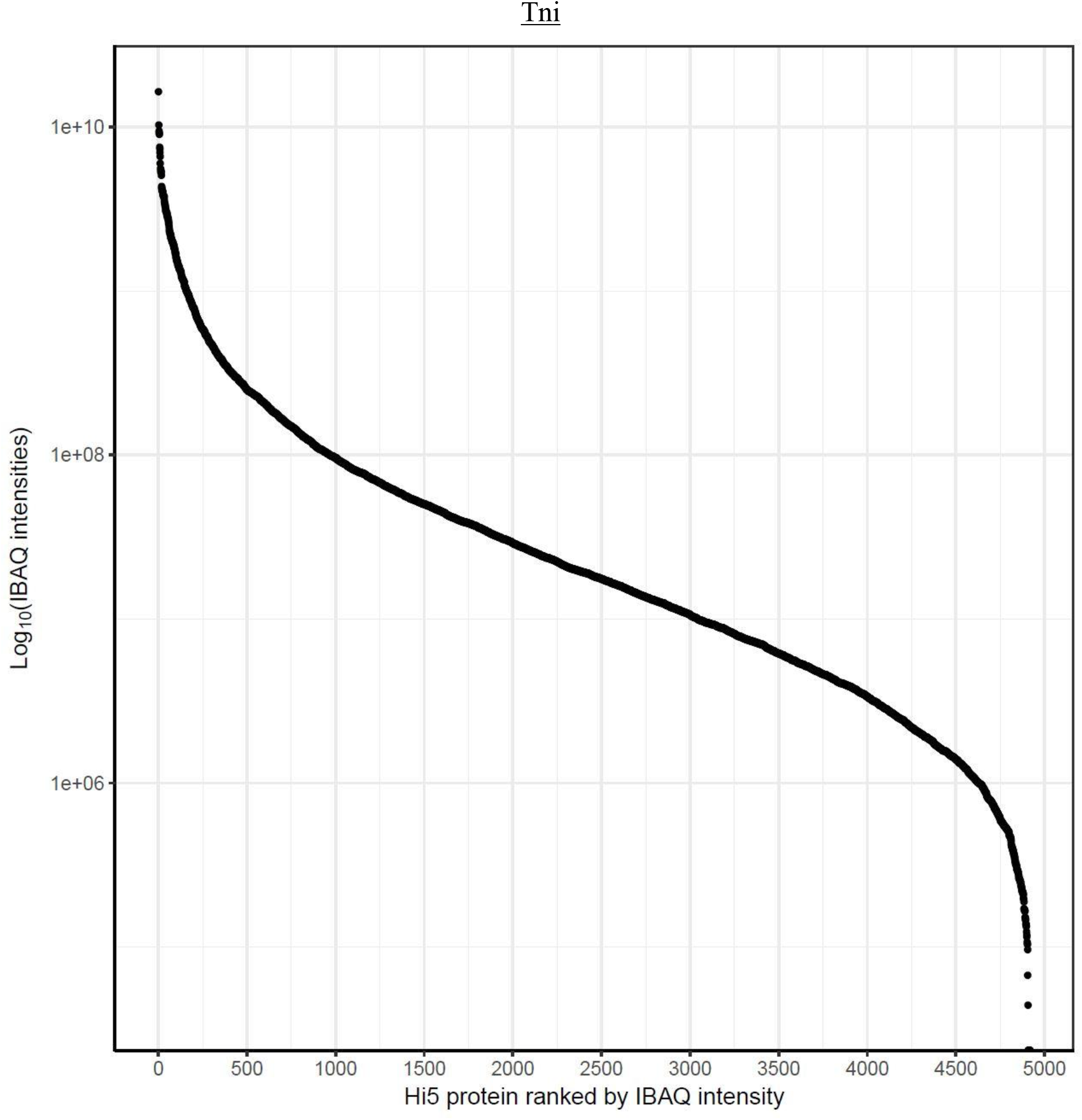
Dynamic range of Sf21 and Tni proteomes. The distribution of Log10(IBAQ intensities) for each protein reveals the typical S-shaped distribution over the 5 orders of dynamic range of MS signals.

Supplementary tables with all information including annotation, expression values and proteomics mapping results can be found in the supplementary files.

### Resource availabilities

The assembled genomes and the corresponding annotation have been deposited in ensembl.lepbase.org. All sequencing reads have been submitted to ENA (European Nucleotide Archive) under the following accession numbers: PRJEB12116/PRJEB24631 (Sf21) and PRJEB24667/PRJEB24632 (Tni). The proteomics dataset is hosted by the ProteomeXchange Consortium and accessible via PRIDE (Vizcaíno et al. 2016) under the following identifier PXD010282. Tutorial for the genome annotation workflow available at https://github.com/galikbence/genome_annotation.

## Discussion

Short-reads assembly remains a challenging problem (Phillippy 2017). The combination of multiple data-types, including mate-pair and long nanopore reads, unlock the possibility to assemble genomes to a useful resolution for research groups with non-model cell lines or organisms. The deep characterization of cell lines enables accurate engineering of its components. The Sf21 genomic information generated in this study, especially the identification of U6 promoter sequences, was already used to improve the introduction of site-specific noncanonical amino acids with unique features for different applications (Koehler et al. 2016). This example underscores the potential applications of this new knowledge as well as the quality of the data generated.

Coupling the sequence information with extensive genome annotation is also crucial for designing meaningful experiments (Bock et al. 2014). The traditional gene annotation procedures are inadequate for these new organisms (Ekblom & Wolf 2014). The publicly available tools provide a good reannotation for well-known organisms, but these tools usually have a limited performance regarding de novo annotation. The workflow presented in this study uses a new combination of publicly available tools as well as external supporting evidence such as transcriptomics information to refine the gene models.

For the first time, proteomes for these 2 cell lines are also characterized and publicly accessible to the community. To date, Bombyx mori was used as a closely related species regarding the protein background landscape. However, using the resources provided with this study, much more reliable and accurate identification of interaction partners of a target protein or potential host cell contaminants, for example, is made feasible.

Cross-species comparison between the two cell lines, Sf21 and Tni, did not reveal any drastic differences on the predicted gene numbers. The functional categories identified at the genomic and proteomic level are also showing a very good correlation between the cells.

The resources provided by this study on Sf21 and Tni cell lines will help the community to better understand these valuable tools and also provide some biological insights on the insects from which these cells were extracted originally: *Spodoptera frugiperda* and *Trichoplusia ni*.

## Materials and Methods

### Cellular culture and nucleic acid extraction

The Sf21 and Tni cells were cultured in Gibco Sf-900 III medium from LifeTechnologies/ThermoFisher and harvested in an early exponential growth stage at 10^6^ cells/ml. 2 ml of culture were centrifuged and a cell pellet containing 2 x10^6^ cells was washed with PBS and then processed for DNA extraction.

Cells or tissues were lysed in 2 PK buffer (200 mM Tris-HCl [pH7.5], 300 mM NaCl, 25 mM EDTA, 2% w/v SDS) containing 200 mg/ml proteinase K at 65°C for 1 hr, extracted with phenol:chloroform:isoamyl alcohol (25:24:1; Sigma, St. Louis, MO), and genomic DNA was collected by ethanol precipitation. The precipitate was dissolved in 10 mM Tris-HCl (pH 8.0), 0.1 mM EDTA, treated with 20 mg/ml RNase A at 37°C for 30 min, extracted with phenol:chloroform:isoamyl alcohol (25:24:1), and collected by ethanol precipitation.

In order to get a global overview of the transcriptome, the cell harvesting procedure was applied at different time points (0h, 4h, 8h and 24h for Tni and 24h and 48h for Sf21). Pellets were then processed for RNA extraction using the RNeasy Kit from Qiagen according to the manufacturer’s instructions.

### DNA sequencing

For each cell line’s DNA, three types of library, (i) two short-insert PE (paired-end) libraries; (ii) two long-insert MP (mate-pair) libraries; (iii) and one TruSeq Synthetic Long-Read library were prepared according to the kit’s specifications and sequenced by Illumina sequencing technology. The reads sequenced from the TruSeq Synthetic Long-Read library were assembled into long synthetic reads larger than 1.5 kb using the TruSeq Long-Read Assembly app v1.1 available on BaseSpace (Illumina Inc.).

Regarding Oxford Nanopore technology (ONT), library preparation was carried out with the Genomic DNA Sequencing Kit SQK-MAP-006 (Oxford Nanopore Technologies). The manufacturer’s instructions were followed, including the NEBNext FFPE DNA repair step (NEB). The final loading mix was assembled with 6 μL pre-sequencing mix, with 4 μL Fuel Mix (Oxford Nanopore Technologies), 75 μL running buffer (Oxford Nanopore) and 66 μL water. DNA sequencing was performed on an r7 flowcell (r7.3) over 48 hours with MinKNOW (V0.51.1.62 and 0.50.2.13, for Sf21 and Tni cells respectively) using the MAP_48Hr_Sequencing_Run_SQK_MAP006.py script. Metrichor™ software was used for basecalling. Poretools performed the conversion from FAST5 to FASTQ formats.

### RNA sequencing

All libraries were prepared using TruSeq Stranded Total RNA (Illumina). Libraries were sequenced by paired-end (Sf21 and Tni) and single-end (Tni) Illumina sequencing technology. All reads have been submitted to ENA (European Nucleotide Archive) under the following accession number: PRJEB24631 (Sf21) and PRJEB24632 (Tni).

### Genome assembly

The global overview of the assembly steps is depicted in Figure 1. At first, the PE reads were corrected and filtered with SGA - version 0.9.43 (Simpson & Durbin 2010). The resulting read pairs were used as input to perform contig assembly, scaffolding and gap closing using SOAPdenovo2 – version 2.4 (Luo et al. 2012).

Secondly, MP reads were processed with FLASH - version 1.2.6 (Magoč & Salzberg 2011) and all overlapping read pairs were discarded. The resulting pairs were employed with SOAPdenovo2 for scaffolding and then gap closing of the previous assembly. Thirdly, the long synthetic (LS) reads were used to scaffold the assembly obtained with PE and MP sequencing data. All data types were then utilized for a gap closing step (SOAPdenovo2).

ONT reads, independently of the 2D quality check, were mapped using the BWA-MEM algorithm (Li 2013) of BWA v0.7.12-r1044 with recommended flags -k11 -W20 -r10 -A1 -B1 -O1 -E1 -L0 -a -Y from the npScarf manual - version 1.6-10a (Cao et al. 2017). The resulting alignment file, as well as the SOAPdenovo2 pre-assembly, were used by npScarf to produce the final assembly.

### Genome assembly quality assessment

The quality of the assembly is assessed by mapping all the PE reads generated to the assembled sequences larger than 10 kb in size using BWA-MEM (Li 2013) aligner (version 0.7.12-r1044). QUAST - version 2.3 (Gurevich et al. 2013) was used to generate table 1 and table S1 with assembly statistics of scaffolds longer than 200bp. The CEGMA - version 2.4 (Parra et al. 2007) benchmark tool was applied to evaluate the completeness of our assemblies by accessing the percentage of ultra-conserved Core Eukaryotic Genes (CEGs) fully and partially present in our dataset.

To compare the newly assembled genome to available data, scaffolds were reordered using MAUVE move contigs function (Darling et al. 2004). Nucmer (MUMmer 3.0) (Kurtz et al. 2004) aligned the sequences of the available genome scaffolds/contigs to our Sf21 genome and the relative position and the orientation are reported.

The largest 28 scaffolds of the Tni genome published by (Fu et al. 2018) were compared to our assembly.

### Transcriptome assembly

RNA content was collected at different time points. All sequencing reads from the different extractions were merged together for further analyses. After quality checking and filtering out duplicates using SeqyClean - version 1.9.10 (Zhbannikov et al. 2013), 77,650,737 and 129,473,322 of RNA-seq (PE and SE) reads for Sf21 and Tni cells respectively, were assembled de novo using Trinity - version 2.0.6 (Grabherr et al. 2011) with default parameters except for in silico read normalisation, where only a subset of reads was used (-- normalize_reads) to increase the assembly quality. Table S2 presents the number of reads kept at each processing step. The assembled transcripts were then cleaned up from duplicates by CD-HIT clustering (Fu et al. 2012), with default minimum similarity of 90%.

### Transcriptome assembly quality assessment

Like the genome quality assessment, the PE and SE reads were mapped back to the assembled transcripts (table S2) using Bowtie2 - version 2.2.6 (Langmead & Salzberg 2012). In addition, the number of full-length protein-coding genes was evaluated using a Trinity tool (analyze_blastPlus_topHit_coverage.pl). Briefly, the assembled transcripts are compared against the SwissProt protein database using BLAST+ - version 2.28 (Camacho et al. 2009). Blast hits are grouped based on similarity to improve the coverage and consolidate the number of full-length protein-coding genes detected. BUSCO (Benchmarking Universal Single-Copy Orthologs) - version 2.0 (Simão et al. 2015) was used to explore the assembly completeness according to conserved orthologs content (insecta_odb9 lineage dataset) such as single-copy, duplicated, fragmented and missing ones across all published and newly assembled transcriptomes. Finally, the assembly stats were computed using one of the Trinity utilities - TrinityStats.pl (Grabherr et al. 2011).

To compare our assembled transcripts to the published data, transcripts from the different sources were mapped back to our assembled genomes using GMAP - version 2016-05-01 (Wu & Watanabe 2005). Additionally, QUAST - version 2.3 (Gurevich et al. 2013) examined the exact similarities between the newly assembled and the previously released transcripts. The metrics were evaluated using our transcripts as a reference (option -R) in both cases. Also, BUSCO was applied to analyze the published datasets using the same parameters as above.

### Genome annotation

The annotation workflow is depicted in figure 5. The following sections describe the different steps of the analysis, which produces gff and fasta files containing all information.

#### Repeat masking

Repeats and other transposable elements (TEs) were identified and masked to increase the gene prediction accuracy and quality. First, these elements were predicted de novo and annotated with RepeatModeler - version 1.0.8 (Smit & Hubley 2010). The generated repeat library was used directly with RepeatMasker - version 4.0.6 (Smit et al. 2015) to identify interspersed repeats and low complexity DNA sequences. The ‘soft’ masking option (-xsmall) was used in order not to confound the gap regions (filled with Ns during scaffolding steps) with the actual repeats in the following step. The used gene prediction tools could handle the soft masked inputs.

#### Gene prediction

In the next step, protein-coding genes were predicted by applying an evidence driven ab initio approach (Yandell & Ence 2012). The general gene prediction algorithm uses predefined gene models and external evidences (transcript and protein information) to find the best gene models. GeneMark-ES - version 4.29 (Ter-Hovhannisyan et al. 2008) and Augustus - version 3.1.0 (Stanke et al. 2004) were employed with different predefined models to predict exon/intron boundaries for protein-coding genes. Three Augustus runs were performed with (i) *H. melpomene* gene model (closest available species), (ii) *H. melpomene* gene model plus the protein sequences derived from the transcriptome assembly as external hints, (iii) the generic gene model plus the protein sequences derived from the transcriptome assembly. The protein sequences were obtained by first predicting open reading frames (ORFs), using a general model in TransDecoder - version 3.0.1 (Haas et al. 2013). Secondly, BLASTP - BLAST+ version 2.2.28+ (Camacho et al. 2009) and HMMER - version 3.1b1 (Finn et al. 2016) searches were achieved on the predicted ORFs using specific releases of SwissProt and Pfam databases provided by Trinity developers. Only sequence hits larger than 100 aa were translated into proteins. This set of protein sequences are used with Augustus as additional evidence to support the best gene model. All other parameters were used with default options. GeneMark-ES was running with default options except for --soft_mask 50 and --evidence.

EvidenceModeler - version 1.1.1 (Haas et al. 2008) was used to combine the different genomic features found by Augustus (Stanke et al. 2004) and GeneMark-ES (Ter-Hovhannisyan et al. 2008) to produce the best consensus gene model. We associated for each of the inputs, the recommended weights according to the EvidenceModeler manual; e.g. 1 for the two ab initio prediction tools, 10 for transcript alignment, 5 for protein alignment and −1 for a repeat. Transcriptome alignment to the assembled genome was obtained using GMAP (Wu & Watanabe 2005) with default parameters. Protein alignment was generated by mapping known Tni and Sf21 protein sequences, taken from NCBI RefSeq database (Release 79), as well as translated protein sequences derived from the transcript assembly (mentioned in previous section) using Scipio - version 1.4.1 (Keller et al. 2008) with default parameters and the outcome from the repeat masking section as repeat information. Gene predictions with less than 3 ab initio support or with no protein or transcript alignment were filtered out. Gene predictions with 3 or more ab initio supports were merged into one and then functionally annotated.

#### Functional annotation

From the consensus gene model generated previously, CDSs were translated into proteins applying standard genetic code. BLASTP was used to align these protein sequences to the insecta subset of NCBI’s non-redundant (nr) protein database (downloaded in October 2016 with insecta taxon; Taxonomy ID: 50557). In order to refine the annotation, proteins returning no hits or mapping to hypothetical proteins were extracted and the corresponding genes were mapped back to the transcriptome. The mapping genes qualified as uncharacterised with in vivo evidence while the rest remain hypothetical.

Protein-domains were predicted by Interproscan - standalone version 5.21-60.0 (Jones et al. 2014), with default parameters against all available databases.

Non-coding RNAs (ncRNA) were identified by Rfam – Infernal - version 1.0.4 and 1.1 (Nawrocki & Eddy 2013) with standard parameters on a reference Rfam database - release 12.0 (Nawrocki et al. 2015). tRNAscan-SE - version 1.3 (Lowe & Eddy 1997) was used with default parameters to identify all transfer RNAs (tRNAs) in the genome. For the ncRNA and the tRNAs identification, both tools used the genome assembly as input.

#### Pathway reconstruction

KEGG Orthologs (KO) were assigned, and pathways were reconstructed using the KEGG Automatic Annotation Server (KAAS) - version 2.1 (Moriya et al. 2007) for “Complete and draft genomes” option. KAAS provides functional gene annotation by comparison against the manually curated KEGG GENES database. Nucleotide sequences, as well as representative Eukaryotes and all insects in the database, were used as input. The bi-directional best hit was applied as the assignment method. The Bioconductor KEGGREST package - version 1.14.1 (Tenenbaum 2018) was used to create coloured reference pathway figures based on Kegg Orthology (KO) IDs.

### Proteomic analysis

#### Sample preparation for Mass Spectrometry

Sf21 or Tni cells were pelleted from 5 mL suspension, to produce a cell pellet containing approximately 10 million cells. The pellets were resuspended in 100 mM Ammonium bicarbonate, pH 7.5 containing Rapigest™ (Waters) at 0.1% to a volume of 1 mL. The cells were lysed by high energy sonication (Bioruptor, Diagenode) and reduced by adding DTT (to a final concentration of 15 mM and heating for 30 minutes at 56 °C), then alkylated using IAA (to a final concentration of 10 mM, 20 minutes in the dark). The proteins were then pelleted using TCA (1 volume ice cold 100% TCA to 4 volumes sample) and left to stand on ice for 30 minutes to precipitate the proteins. The samples were then centrifuged at 14,000 rpm for 20 minutes, 4 °C. After removal of the supernatant, the precipitates were washed once with 1000 μL 10% TCA, vortexed, centrifuged again for 20 minutes at 4°C, then washed twice (2 × 1,000 μL with ice cold (stored at −20 °C before use) acetone. Vortexing and centrifugation steps were repeated as before. The pellets were then allowed to air-dry before being dissolved by sonication in digestion buffer (1 mL, 4M urea in 0,1M HEPES, pH 8); To a 50 μL aliquot, estimated to contain around 50-100 μg protein, 1 μg LysC enzyme (Wako) was added and incubated for 4h at 37 °C. Then the samples were diluted with milliQ water (to reach 1.5M urea) and were incubated with 1 μg trypsin for 16 h at 37 °C. The digests were then acidified with 10% trifluoroacetic acid and then desalted with Waters Oasis^®^ HLB μElution Plate 30μm in the presence of a slow vacuum. In this process, the columns were conditioned with 3 × 100 μL solvent B (80% acetonitrile; 0.05% formic acid) and equilibrated with 3 × 100 μL solvent A (0.05% formic acid in milliQ water). The samples were loaded, washed 3 times with 100 μL solvent A, and then eluted into PCR tubes with 50 μL solvent B. The eluates were dried down with the speed vacuum centrifuge and dissolved in 50 μL 5% acetonitrile, 95% milliQ water, with 0.1% formic acid prior to pre-fractionation by high pH reverse phase chromatography (HpH) to increase proteome coverage.

#### Offline high pH fractionation

Offline high pH reverse phase fractionation was performed using an Agilent 1260 (Tni) or an Agilent 1200 (Sf21) Infinity HPLC System equipped with a binary (Tni) or quarternary (Sf21) pump, degasser, variable wavelength UV detector (set to 220 and 254 nm), peltier-cooled autosampler (set at 10°C) and a fraction collector. The column was a Waters XBridge C18 column (3.5 μm, 100 × 1.0 mm, Waters) with a Gemini C18, 4 × 2.0 mm SecurityGuard (Phenomenex) cartridge as a guard column. The solvent system consisted of 20 mM ammonium formate (pH 10.0) as mobile phase (A) and 100% acetonitrile as mobile phase (B). 50 μg of the resuspended peptides after digestion and clean up (dissolved in 5% ACN, water, 0.1% FA) were injected, and the separation was accomplished at a mobile phase flow rate of 0.1 mL/min using a linear gradient from 100% A to 31 % B in 91 min. 48 fractions were collected along with the LC separation, which were subsequently pooled into 10 discordant fractions. Pooled fractions were dried in a Speed-Vac and then stored at −80°C until LC-MS/MS analysis, where approximately 1 μg per fraction were used to acquire DDA data.

#### Sf21 Data Acquisition

Peptides were separated using the Dionex Ultimate 3000 nUPLC system (Thermo) fitted with a trapping cartridge (Acclaim PepMap C18, 5 μm, 300μm × 5 mm) and an analytical column (PepMap C18, 1.8 μm, 75 μm × 500 mm). The outlet of the analytical column was coupled directly to a Q-Exactive+ (Thermo Fisher Scientific) using the Proxeon nanospray source. Solvent A was water, 0.1% (v/v) formic acid and solvent B was acetonitrile, 0.1% (v/v) formic acid. The samples were loaded with a constant flow of 99% solvent A at 30 μL/min, onto the trapping column. Trapping time was 4 min. Peptides were eluted via the analytical column at a constant flow of 0.3 μL/min, with both trap and analytical columns being held at 40 °C. During the elution step, the percentage of solvent B increased in a linear fashion after isocratic flow at 2% B for 4 minutes, to 7% B in 2 minutes, then from 7% B to 24% B in a further 80 min and to 40% B by 115 min. The peptides were introduced into the mass spectrometer via a Pico-Tip Emitter 360 μm OD × 20 μm ID; 10 μm tip (New Objective) and a spray voltage of 2.2kV was applied. The capillary temperature was set at 300 °C. Full scan MS spectra with mass range 350-1650 m/z were acquired in profile mode in the 70,000. The filling time was set at maximum of 32 ms with an AGC target of 1 × 10^6^ ions. The peptide match algorithm was disabled and only charge states from 2^+^ to 5^+^ were selected for fragmentation. The top 10 most intense ions from the full scan MS were selected for MS2, using quadrupole isolation and a window of 2 Da. An intensity threshold of 7 x10^4^ ions was applied. HCD was performed with collision energy of 30%. A maximum fill time of 65 ms, with an AGC target of 5 × 10^5^ for each precursor ion was set. MS2 data were acquired in centroid with a resolution of 17,500 with fixed first mass of 110 m/z. The dynamic exclusion list was with a maximum retention period of 25 sec and relative mass window of 10 ppm. Isotopes were also excluded.

#### Tni Data Acquisition

Peptides were separated using the nanoAcquity UPLC system (Waters) fitted with a trapping (nanoAcquity Symmetry C18, 5 μm, 180 μm × 20 mm) and an analytical column (nanoAcquity BEH C18, 2.5 μm, 75 μm × 250 mm). The outlet of the analytical column was coupled directly to an Orbitrap Fusion Lumos (Thermo Fisher Scientific) using the Proxeon nanospray source. Solvent A was water, 0.1% (v/v) formic acid and solvent B was acetonitrile, 0.1% (v/v) formic acid. The samples were loaded with a constant flow of solvent A at 5 μL/min, onto the trapping column. Trapping time was 6 min. Peptides were eluted via the analytical column at a constant flow of 0.3 μL/min, at 40°C. During the elution step, the percentage of solvent B increased in a linear fashion from 5% to 7% in 10 minutes, then from 7% B to 30% B in a further 105 min and to 45% B by 130 min. The peptides were introduced into the mass spectrometer via a PicoTip Emitter 360 μm OD × 20 μm ID; 10 μm tip (New Objective) and a spray voltage of 2.2kV was applied. The capillary temperature was set at 300°C. Full scan MS spectra with mass range 375-1500 m/z were acquired in profile mode in the Orbitrap with a resolution of 60,000 using the quad isolation. The RF on the ion funnel was set to 30%. The filling time was set at maximum of 50 ms with an AGC target of 2 × 10^5^ ions and 1 microscan. The peptide monoisotopic precursor selection was enabled and only charge states from 2^+^ to 7^+^ were selected for fragmentation. The most intense ions (instrument operated for a 3 second cycle time) from the full scan MS were selected for MS2, using quadrupole isolation and a window of 1.4 Da. An intensity threshold of 5 x10^3^ ions was applied. HCD was performed with collision energy of 30%. A maximum fill time of 22 ms with an AGC target of 1 × 10^5^ for each precursor ion was set. MS2 data were acquired in centroid in the Orbitrap with a resolution of 15,000, with fixed first mass of 120 m/z. The dynamic exclusion list was with a maximum retention period of 15 sec and relative mass window of 10 ppm. Isotopes were also excluded.

#### MS Data Processing

Using protein fasta files, generated from the translated assembled transcripts obtained from the genome annotation pipeline, (for the Sf21 and Tni data 10 fractions separately) were searched with MaxQuant (v.1.5.3.28) and the Andromeda search engine (Cox & Mann 2008). The data in both cases were searched against the generated species specific (Sf21 or Tni) database with a list of common contaminants appended. The data were searched with the following modifications: Carbamidomethyl (C) (Fixed) and Oxidation (M) and Acetyl (Protein N-term) (Variable). The mass error tolerance for the full scan MS spectra was set at 20 ppm and for the MS/MS spectra at 0.01 Da. A maximum of 2 missed cleavages were allowed. From the MaxQuant searches, iBAQ (intensity-based absolute quantification) values were exported and used to generate figure 8.

## Acknowledgements

The authors thank the P4EU network (https://p4eu.org) for providing resources necessary for this study. The FLI is a member of the Leibniz Association and is financially supported by the Federal Government of Germany and the State of Thuringia. The authors gratefully acknowledge support from the FLI proteomics core facility. The authors would also like to thank the Genomics and the Proteomics Core Facility at EMBL. The Vienna Biocenter Core Facilities (VBCF) receives funding from the City of Vienna and the Austrian Federal Ministry of Education, Science, and Research. PSB acknowledges funding from the Laura Bassi Centres of Expertise initiative for the Centre of Optimized Structural Studies, project 253275.

## Author contribution

Conceptualization, P.S., A.G. and V.B.; Methodology, B.G., J.J.M.L., J.M.K., M.H.F., B.B., J.B., B.H., P.G.C., R.H., D.P., P.S., H.B., K.R., A.G. and V.B.; Validation, B.G., J.J.M.L., J.M.K., M.H.F., B.B., J.B., B.H., P.G.C., R.H., D.P., P.S., K.R., A.G. and V.B.; Formal analysis, B.G., J.J.M.L., J.M.K., P.S., A.G. and V.B.; Investigation, B.G., J.J.M.L., J.M.K., M.H.F., B.B., J.B., B.H., P.G.C., R.H., D.P., P.S., H.B., K.R., A.G. and V.B.; Resources P.S., A.G. and V.B.; Data curation, J.J.M.L and J.M.K.; Writing—original draft preparation, B.G., J.J.M.L., M.H.F., B.B., J.B., B.H., P.G.C., R.H., D.P., P.S., H.B., K.R., A.G. and V.B.; Writing—review and editing, B.G., J.J.M.L., J.M.K., M.H.F., B.B., J.B., B.H., P.G.C., R.H., D.P., P.S., H.B., K.R., A.G. and V.B.; Visualization, B.G., J.J.M.L.; Supervision, P.S., A.G. and V.B.; Project administration, P.S., A.G. and V.B.; Funding acquisition, B.G., P.S., A.G and V.B.

