## Supplementary tables and figures for "Extensive OMICS resource for Sf21 and Tni cell lines"

| | Contigs assembly only with PE reads | Assembly with PE and MP reads | Assembly with PE, MP and SLR reads | Assembly with PE, MP, SLR and ONT reads |
|---|---|---|---|---|
| **Sf21 genome** | | | | |
| # sequences (>= 1 Kb) | 47,866 | 17,588 | 11,871 | 4,020 |
| Largest scaffold (bp) | 264,592 | 1,209,752 | 2,608,367 | 2,900,257 |
| Total length (bp) | 430,604,418 | 477,599,629 | 488,468,751 | 463,041,686 |
| N50 (bp) | 18,101 | 140,626 | 307,038 | 364,523 |
| L50 | 6,168 | 843 | 383 | 315 |
| GC (%) | 36.20 | 36.33 | 38.59 | 38.42 |
| # N's per 100 Kb | 27.82 | 3559.42 | 776.82 | 681.43 |
| **Tni genome** | | | | |
| # sequences (>= 1 Kb) | 53,137 | 22,485 | 14,683 | 2,954 |
| Largest scaffold (bp) | 101,314 | 595,205 | 1,711,767 | 1,722,548 |
| Total length (bp) | 355,056,269 | 393,087,391 | 407,374,447 | 332,103,479 |
| N50 (bp) | 8,666 | 82,532 | 252,810 | 326,309 |
| L50 | 9,637 | 1,114 | 395 | 293 |
| GC (%) | 36.08 | 35.82 | 36.51 | 36.30 |
| Number of N's per 100 Kb | 57.75 | 1183.12 | 575.45 | 499.33 |

**Table S1: Short summary of Sf21 and Tni *de novo* genome assemblies using progressively different types of reads.** The N50 value is defined as the sequence length of the shortest contig at 50% of the total genome length. L50 value represent the smallest number of contigs to which their cumulative length represents half of genome size.

| | Sf21 | Tni |
|---|---|---|
| | **RNA-Seq data** | |
| Total number of PE/SE reads | 91,741,494 | 240,178,919 |
| Number of PE/SE reads after adaptor and quality trimming | 86,754,674 | 164,941,193 |
| Number of PE/SE reads after duplicate removal | 77,650,737 (#PE: 75,773,152 #SE:1,877,585) | 129,473,322 (#PE: 124,582,705 #SE: 4,890,617) |
| | **Transcriptome assembly** | |
| Number of assembled transcripts | 28,339 | 57,600 |
| Number of assembled transcripts (>= 1000 bp) | 9,948 | 21,534 |
| GC (%) | 41.31 | 42.93 |
| Total assembled bases | 31,697,415 | 67,831,071 |
| N50 (bp) | 1,956 | 2,076 |
| Median transcript length (bp) | 611 | 648 |
| Mean transcript length (bp) | 1,118.51 | 1,177.65 |
| Largest transcript (bp) | 24,940 | 20,156 |
| # Full length transcripts (>= 80%) | 3,211 | 3,669 |
| Percentage of mapped RNAseq PE/SE reads | ~ 97% (proper pairs) | ~ 69% (proper pairs) |
| Number of transcripts after CD-HIT run (90% similarity) | 24,992 | 41,041 |

**Table S2: Sf21 and Tni *de novo* transcriptome sequencing data and assembly features.** This table presents the total number of sequencing paired-end (PE) reads and the number of processed reads after trimming and duplicate removal. The filtered reads were then used for transcriptome assembly. The shown metrics cover the number of assembled transcripts, the GC percent, the cumulative length of the assembly, the N50, the median, the mean and the maximum length of the transcripts, the full-length transcript coverage and the number of RNAseq PE reads mapping back to the assembled sequences. Finally, the assembled transcripts were clustered, before the gene prediction step, using CD-HIT to remove duplications and clean up the dataset.

| | **Sf21** | **Tni** |
|---|---|---|
| | **Main features** | |
| Number of tRNAs | 1,233 (311 pseudos) | 1,965 (1,166 pseudos) |
| Number of ncRNAs | 396 | 249 |
| Number of repeats (excluding low complexity regions and satellites) | 431,477 | 390,271 |
| Number of protein-coding genes | 21,506 | 14,159 |
| | **Exon statistics** | |
| Exon count | 110,027 | 89,378 |
| Average number of exons/gene | ~5.12 | ~6.31 |
| Exon space count (bp) | 30,421,075 | 20,631,338 |
| Average exon size (bp) | 276 | 231 |
| Median exon size (bp) | 160 | 156 |
| Minimum exon size (bp) | 14,440 | 14,890 |
| Maximum exon size (bp) | 3 | 3 |
| | **Intron statistics** | |
| Intron count | 88,521 | 75,220 |
| Number of introns/gene | ~4.12 | ~5.31 |
| Intron space count (bp) | 94,576,984 | 77,902,387 |
| Average intron size (bp) | 1086 | 1036 |

| | | |
|---|---|---|
| Median intron size (bp) | 649 | 597 |
| Minimum intron size (bp) | 86,400 | 79,230 |
| Maximum intron size (bp) | 21 | 22 |
| **Intergenic space statistics** | | |
| Intergenic space count | 13,035 | 8,735 |
| Intergenic space size (bp) | 353,235,779 | 248,980,732 |
| Average intergenic space distance (bp) | 31,190 | 33,600 |
| Median intergenic space distance (bp) | 12,840 | 15,390 |
| Minimum intergenic space distance (bp) | 1 | 1 |
| Maximum intergenic space distance (bp) | 977,500 | 584,000 |

**Table S3. Genomics feature count.** This table represents the genomics feature characteristics and statistics for the Sf21 and Tni genomes.

### Supplementary figures

Sf21

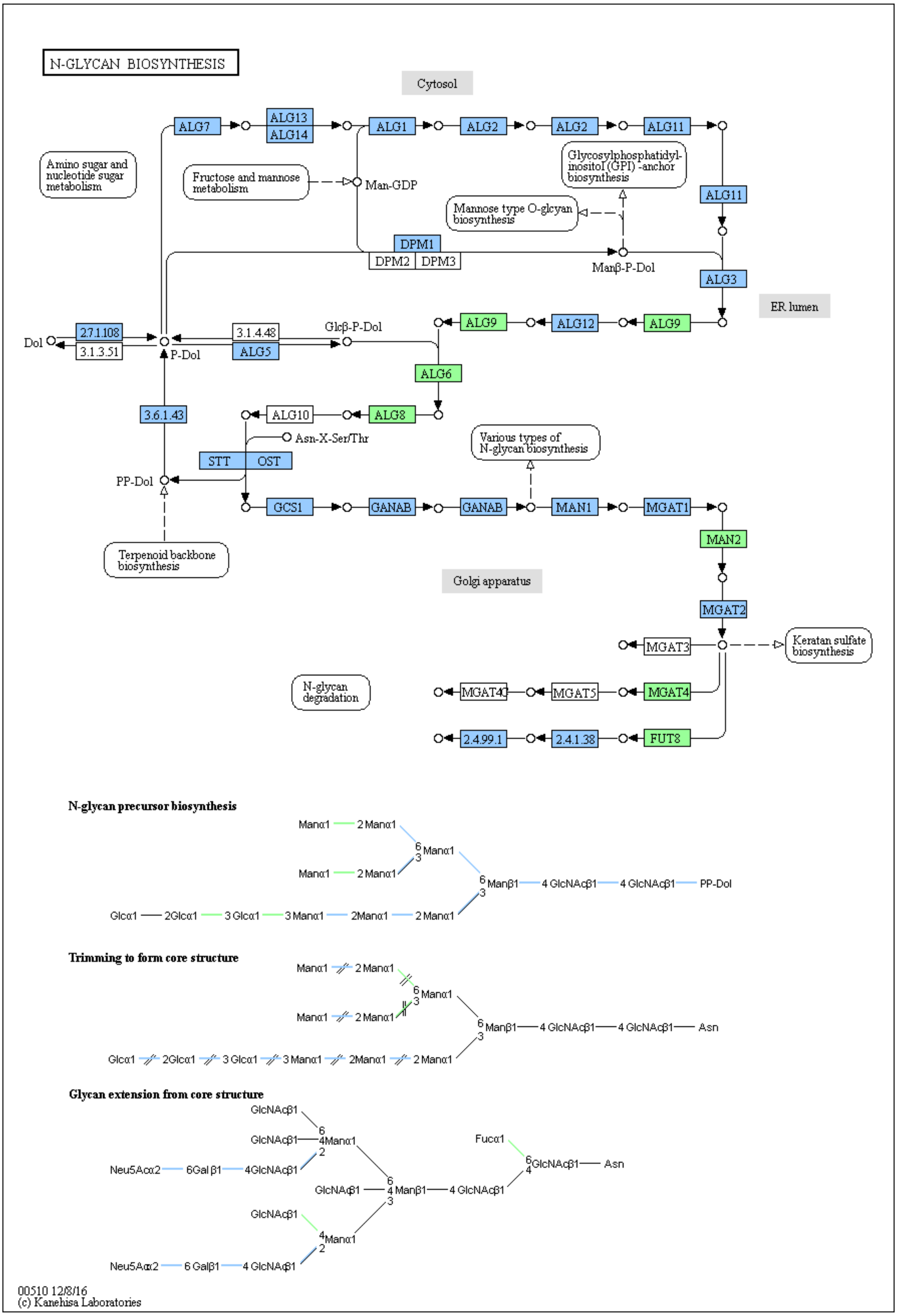

Tni

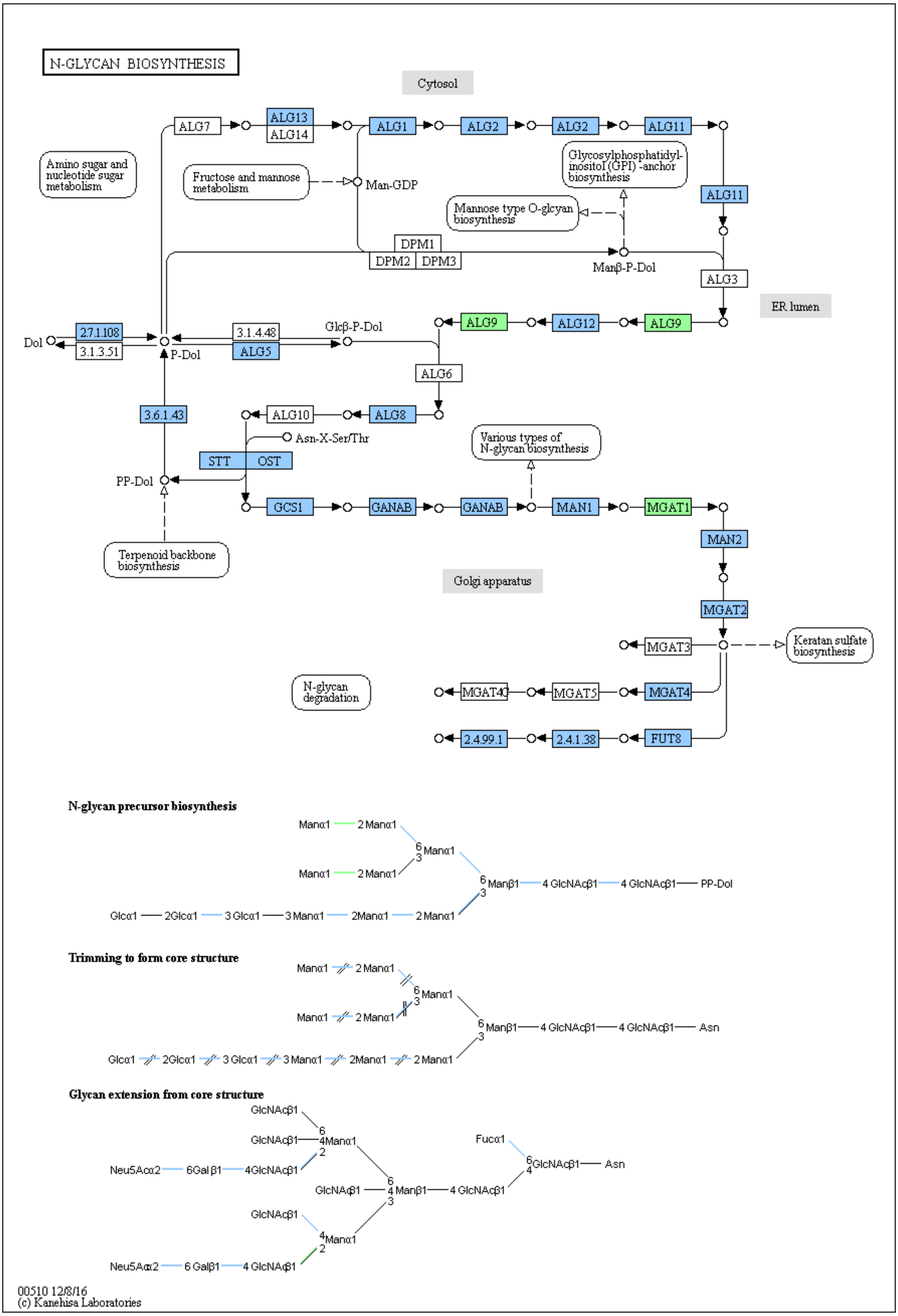
N-GLYCAN BIOSYNTHESIS
Cytosol
ALG7
ALG13
ALG14
ALG1
ALG2
ALG2
ALG11
Amino sugar and nucleotide sugar metabolism
Fructose and mannose metabolism
Man-GDP
Glycosylphosphatidyl-inositol (GPI) -anchor biosynthesis
ALG11
Mannose type O-glcyan biosynthesis
DPM1
DPM2
DPM3
Manβ-P-Dol
ALG3
ER lumen
Dol
2.7.1.108
3.1.3.51
P-Dol
3.1.4.48
ALG5
Glcβ-P-Dol
ALG9
ALG12
ALG9
ALG6
3.6.1.43
ALG10
ALG8
Asn-X-Ser/Thr
STT
OST
Various types of N-glycan biosynthesis
PP-Dol
GCS1
GANAB
GANAB
MAN1
MGAT1
MAN2
Terpenoid backbone biosynthesis
Golgi apparatus
MGAT2
MGAT3
Keratan sulfate biosynthesis
N-glycan degradation
MGAT4C
MGAT5
MGAT4
2.4.99.1
2.4.1.38
FUT8
N-glycan precursor biosynthesis
Manα1
2 Manα1
6 Manα1 3
Manα1
2 Manα1
6 Manβ1
4 GlcNAcβ1
4 GlcNAcβ1
PP-Dol
3
Glcα1
2Glcα1
3 Glcα1
3 Manα1
2Manα1
2 Manα1
Trimming to form core structure
Manα1
2 Manα1
6 Manα1 3
Manα1
2 Manα1
6 Manβ1
4 GlcNAcβ1
4 GlcNAcβ1
Asn
3
Glcα1
2Glcα1
3 Glcα1
3 Manα1
2Manα1
2 Manα1
Glycan extension from core structure
GlcNAcβ1
6
GlcNAcβ1
4Manα1
2
Neu5Acα2
6Gal β1
4GlcNAcβ1
Fucα1
6 GlcNAcβ1
Asn
4
GlcNAcβ1
6
4 Manβ1
4 GlcNAcβ1
3
GlcNAcβ1
4 Manα1
2
Neu5Acα2
6 Galβ1
4 GlcNAcβ1
00510 12/8/16
(c) Kanehisa Laboratories

*B. mori*

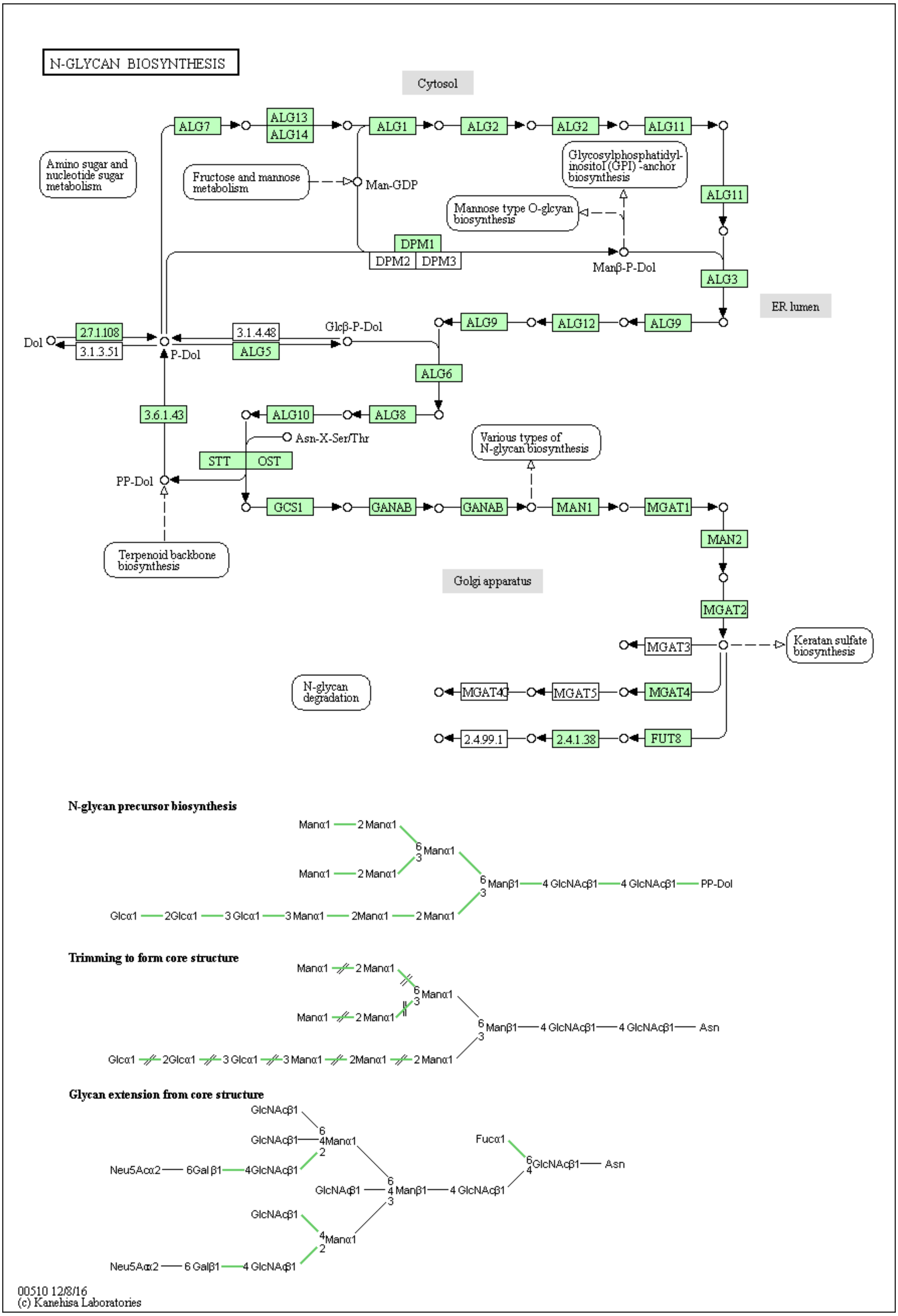
N-GLYCAN BIOSYNTHESIS
Cytosol
ER lumen
Golgi apparatus
ALG7
ALG13
ALG14
ALG1
ALG2
ALG11
Amino sugar and nucleotide sugar metabolism
Fructose and mannose metabolism
Man-GDP
Glycosylphosphatidyl-inositol (GPI) -anchor biosynthesis
Mannose type O-glcyan biosynthesis
DPM1
DPM2
DPM3
Manβ-P-Dol
ALG3
Dol
2.7.1.108
3.1.3.51
P-Dol
3.1.4.48
ALG5
Glcβ-P-Dol
ALG9
ALG12
ALG6
3.6.1.43
ALG10
ALG8
Asn-X-Ser/Thr
STT
OST
PP-Dol
Various types of N-glycan biosynthesis
GCS1
GANAB
MAN1
MGAT1
MAN2
Terpenoid backbone biosynthesis
MGAT2
MGAT3
Keratan sulfate biosynthesis
N-glycan degradation
MGAT4C
MGAT5
MGAT4
2.4.99.1
2.4.1.38
FUT8
N-glycan precursor biosynthesis
Trimming to form core structure
Glycan extension from core structure
Manα1
2 Manα1
Manα1
Manβ1
4 GlcNAcβ1
PP-Dol
Glcα1
2Glcα1
3 Glcα1
3 Manα1
2Manα1
Asn
GlcNAcβ1
4Manα1
Neu5Acα2
6Gal β1
4GlcNAcβ1
Fucα1
GlcNAcβ1
4 Manβ1
4 GlcNAcβ1
6Galβ1
00510 12/8/16
(c) Kanehisa Laboratories

**Figure S1. N-glycan pathway.** N-glycan pathway representations for SF21 and Tni, as well as Bombyx Mori. Predicted genes, based on the genomic analysis, are marked with green and predicted and expressed, based on the genomic and transcriptomic analysis, are represented in blue.
